# Evidence for HARKing in mouse behavioural tests of anxiety

**DOI:** 10.1101/2022.12.01.518668

**Authors:** Marianna Rosso, Adrian Herrera, Hanno Würbel, Bernhard Voelkl

## Abstract

Anxiety disorders are among the most common mental health conditions needing new and better treatments. Research and development of anxiolytic drugs using animal models heavily rely on behavioural tests, most of which measure anxiolytic effects by increased exploratory activity in potentially threatening environments. However, interpretation of such tests is ambiguous because reduced exploratory activity can reflect either anxiety or sedation by the compound. Based on a systematic review of the sensitivity of mouse behavioural tests of anxiety to anxiolytic drugs, we analysed outcomes of 206 studies testing the effect of diazepam in either the open-field test or the hole-board test, as outcomes of these tests showed substantial variation in both direction and strength of the drug effect. Surprisingly, we found that both the rationale given for using the test, whether to detect anxiolytic or sedative effects, and the predicted effect of diazepam, anxiolytic or sedative, strongly depended on the reported test results. The most likely explanation for such strong dependency is post-hoc reasoning, also called hypothesizing after the results are known (HARKing). HARKing can invalidate study outcomes and hampers evidence synthesis by inflating effect sizes. It may also lead researchers into blind alleys, put patients in clinical trials at risk, and waste animals, time, and resources for inconclusive research.

## Main text

Recently, the validity of behavioural tests for anxiety in rodents has been questioned, as they often fail to replicate results with different anxiolytic compounds or different mouse models (Bespalov and Steckler 2021; Ennaceur and Chazot 2016). To investigate whether this critique is justified, we have previously conducted a systematic review on the sensitivity of a wide range of behavioural test measures of common mouse behavioural tests of anxiety to detect anxiolytic drugs approved for treating anxiety in humans (Rosso et al. 2022). Across the 2476 outcomes analysed, we had found considerable variation in both direction and magnitude of effects within the same test measure, compound, and dosage. Although some variation in effect size between studies is expected, we found contradictory results. Notably, for two test measures— “number of squares crossed” in the Open Field Test and “number of head dips” in the Holeboard Test—, the most frequently tested compound, diazepam produced almost equal numbers of positive and negative effects despite comparable test conditions. To understand these contradicting results, we further analysed this pool of research papers by assessing the interpretation of the test results and its relation to the rationales given for performing these tests, and the predicted effects of the compound.

The analysis comprised 241 outcomes from 206 research publications assessing the effect of diazepam, of which 151 outcomes (from 149 publications) were based on “number of squares crossed” in the Open field test and 90 (from 90 publications) were based on “number of head dips” in the Holeboard test. A maximum dosage of 2 mg/kg diazepam was chosen as inclusion criterion to ensure that results were comparable.

For each paper, two independent reviewers extracted information regarding (i) the interpretation of the observed effect of diazepam, (ii) the test rationale, and (iii) the predicted effect of diazepam. This was done by searching for statements which would answer the following three questions. First, were the observed behavioural changes elicited by diazepam described as sedative or anxiolytic effects? Second, was the test measure according to the authors meant to measure anxiety or sedative effects? Third, was diazepam administered as a sedative drug or as an anxiolytic drug?

While in some cases the study authors explicitly addressed these questions, it also happened frequently that the authors either did not elaborate on these questions or made vague or ambiguous statements (e.g., the authors wrote that the test measure was used to “assess anxiety, locomotion and emotionality”). In those cases, we classified the authors’ stance as ‘not reported or ambiguous’.

In both tests, a decrease in activity consistently led researchers to conclude that diazepam had a sedative or depressant effect, while an increase in activity was almost always interpreted as diazepam having an anxiolytic effect (Fig 1,2). This result fits with our expectation based on the description of these tests by their inventors (Hall 1934; File and Wardill 1975). However, when it comes to the predicted effect of diazepam and the test rationale, we observed clear discrepancies in the study descriptions.

**Figure 1:**
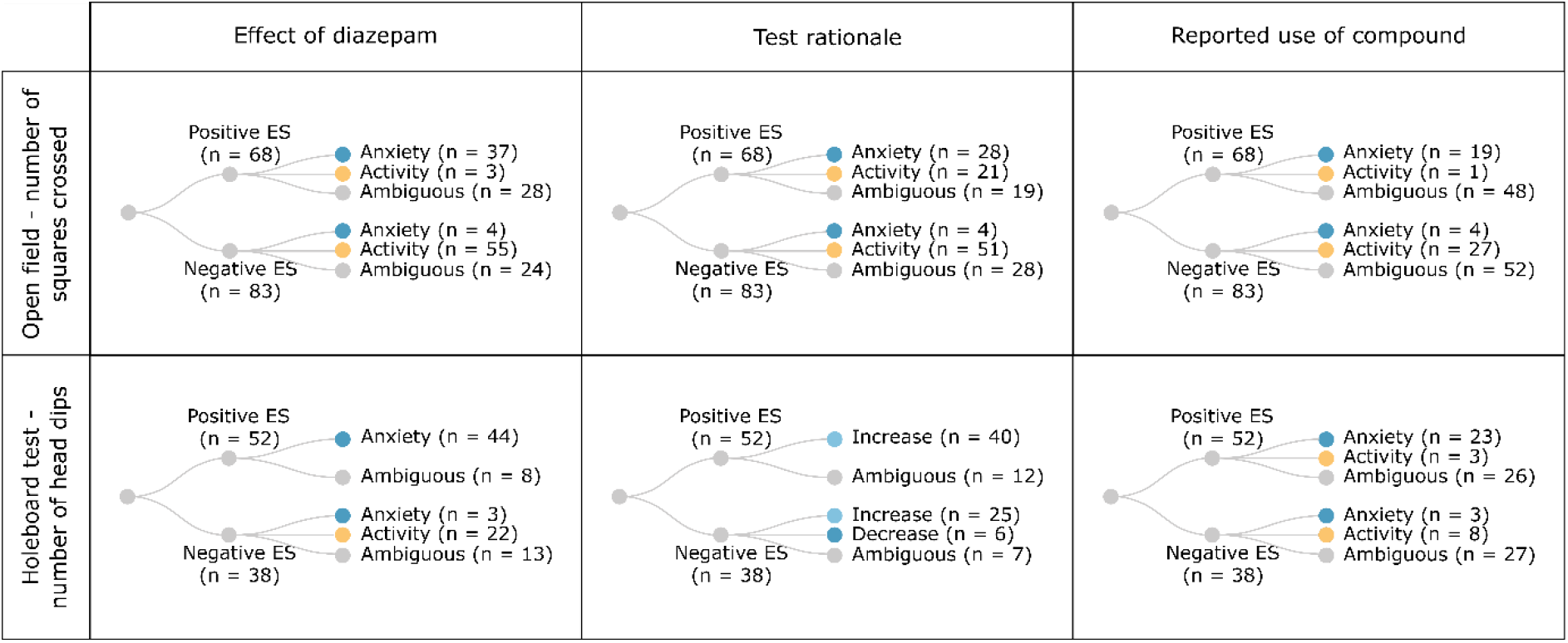
Tree diagram displaying the distribution of outcomes across outcome measure, effect size (ES), item of analysis, and reporting.

**Figure 2:**
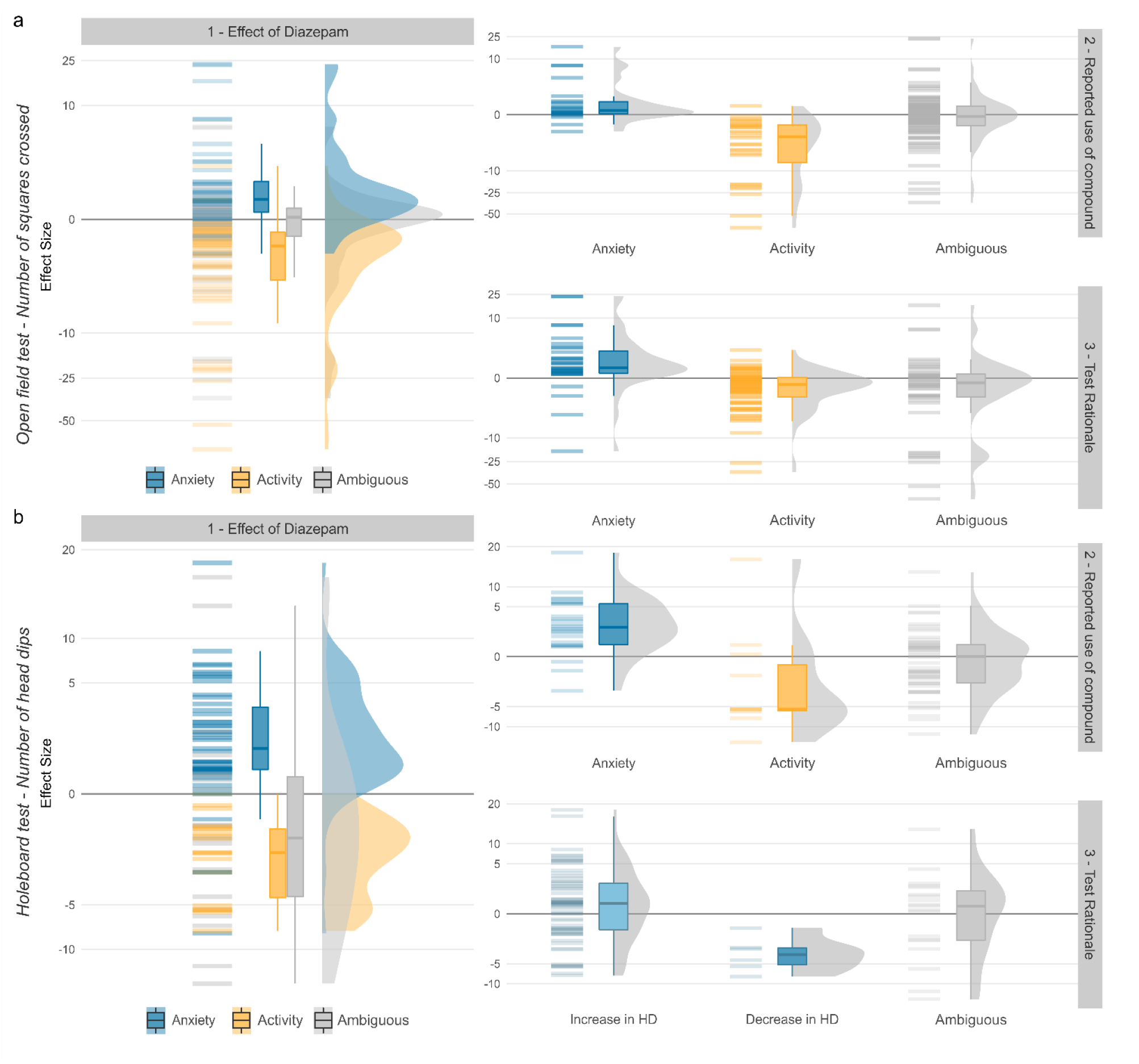
Relation between 1) effect of diazepam, 2) reported use of diazepam, and 3) rationale for the use of the test measure and effect size(y-axis), for number of squares crossed in the Open field test (a) and number of head dips in the Holeboard test (b). Horizontal bars, boxplots, and violin plots indicate the distribution of the effect sizes as extracted from the publications.

The reported rationale for the use of diazepam showed an unexpected bi-directional pattern. In 42 out of 46 cases where a positive effect size was detected and a prediction for the effect of diazepam was made (both tests combined), researchers described diazepam as a standard anxiolytic drug. However, when researchers found a negative effect size, diazepam was described as a standard sedative drug in 35 out of the 42 cases in which a prediction was made. In 79 cases, where the authors did not make a prediction for the effect of diazepam, or the explanation was ambiguous, both negative and positive effect sizes were reported approximately equally often, and the summary effect size was very close to zero (Fig 1,2).

Similarly, the rationale provided for measuring ‘numbers of squares crossed’ in the Open field test, was dependent on the study outcome. In 28 out of 49 cases where a positive effect was detected and a rationale was given, researchers presented the test measure as a tool to assess anxiety, while in 51 out of 55 studies where a negative effect was detected and a rationale was given, researchers described the same test measure as assessment tool for locomotor activity. In the absence of a directional prediction about the effect of diazepam or an unambiguous rationale we found both negative and positive effect sizes approximately equally often, and the summary effect size was very close to zero.

In all studies, researchers used comparable dosages (ranging overall from 0.1 mg/kg to 2 mg/kg, whereby in 120 studies researchers used the same dosage of 1 mg/kg) and usually the same method and route of administration. Therefore, we should assume that the observed variation in reported effect sizes is primarily due to random sample variation. As such, it should be independent of the researchers’ predictions about the effect of the compound or their rationale for using the test measure. This is, however, not the case. There are two possible explanations for these unexpected results – publication bias and HARKing. Thus, such deviation from random variation could be explained, if researchers were more likely to publish results that matched their predictions (the file drawer problem, Rosenthal, 1979), or, alternatively, if they formulated the test rationale or the predicted effect of the compound after the results of the behavioural tests had been known.

Given these two alternative explanations, the question arises how strong a tendency for HARKing or publication bias, or a combination of both, would have to be, to explain the discrepancies in the test rationales and predicted effects discovered in the present meta-analysis. Based on the reported effects, we can calculate expected outcomes for each combination of researchers openly reporting their results, researchers tweaking their explanations based on outcomes (HARKing) and researchers suppressing results contrary to their expectations (publication bias). For the predicted effect, i.e., whether diazepam was declared to have an anxiolytic or sedative effect, Fig. 3a suggests that in the absence of publication bias, the observed reporting pattern for numbers of squares crossed in the open field test are best described if we assume that 80% of researchers were HARKing. Assuming a publication bias of e.g., 40%, the observed pattern still suggests 68% of HARKing among those papers finally published. For the number of head dips in the hole board test, reporting patterns suggests a prevalence of HARKing of 64% in the absence of bias, and still 45% with 40% publication bias (Fig 3c). Similarly, for the rationale of the test, i.e., whether a low number of squares crossed in the open field test indicates anxiety or a sedative effect of the drug, we found that—in the absence of publication bias—a prevalence of 51% HARKing and—for a publication bias of 40%—32 percent of HARKing would be the most likely explanation for the observed reporting pattern (Fig 3b).

**Figure 3:**
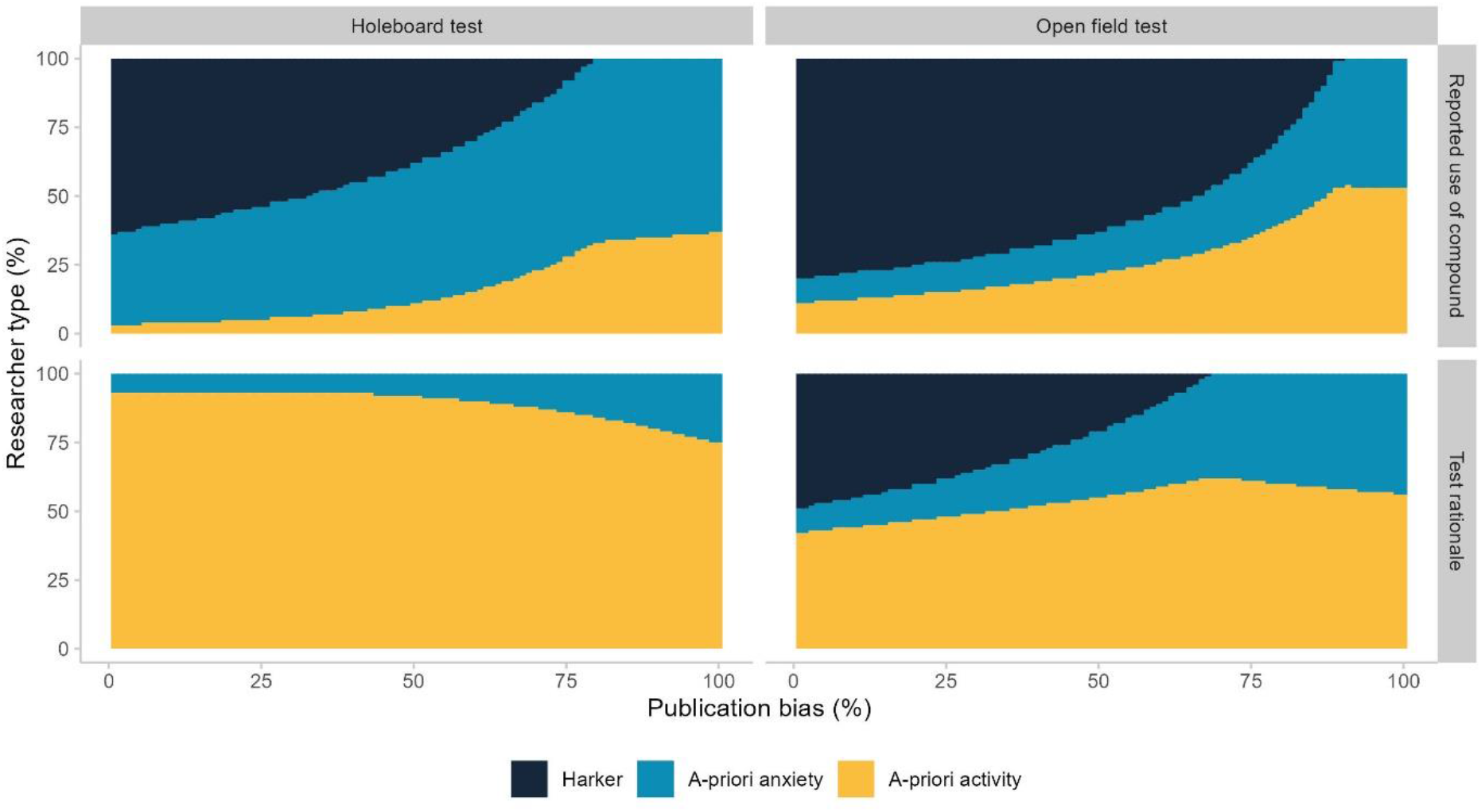
Most likely composition of the population of researchers given the percentage of publication bias. Light blue: researchers who declare their belief that diazepam has an anxiolytic effect and that the test measure is a measure of anxiety before data analysis; Yellow: researchers who declare their believe that diazepam has a sedative effect and that the test measure is a measure of locomotor activity before data analysis; Dark blue: researchers that define the expected effect of the compound and the rationale of the test after data analysis and in accordance with the outcome (HARKers). (a) Rationale behind the use of compound in the OF test, (b) rationale for the test measure (number of squares crossed) in the OF test, (c) rationale behind the use of compound in the HB test, (d) rationale for the test measure in the HB test (number of head dips).

Using behavioural measures of anxiety as an example, we identify a severe problem with current standards of conducting and reporting animal studies. The ambiguity in the test rationale and drug effect together with the flexibility in the interpretation of the drug effects seem to hinder critical evaluation of the validity and replicability of these test measures. If all researchers had agreed that the number of squares crossed in the open field test is a measure of anxiety, then the authors of 66 out of 151 studies would have reported that diazepam did not have an anxiolytic (and in many cases had an anxiogenic) effect, instigating a discussion about the reliability and validity of that test measure. If, on the other hand, all researchers had agreed that the number of squares crossed is simply a measure of locomotor activity to detect sedative effects of a compound, the remaining 85 researchers would have reported that diazepam did not have a sedative effect, again questioning the reliability or validity of that test measure. However, we found that not in a single one out of the 206 studies, authors mentioned that the observed effects of diazepam on the respective test measure were against their expectations.

HARKing represents a major threat to a hypothesis-testing framework, where studies are designed to falsify (null) hypotheses (Kerr 1998). As a form of analytical flexibility, it undermines the robustness and reproducibility of published research for several reasons (Munafò et al. 2017; Bishop 2019). It hinders the process of falsification of false hypotheses, while inflating type I errors, that is, false-positive results mistaken for true effects. It generates a bias in the published literature, missing null results (Wicherts et al. 2016; Rubin 2017, 2020; Nosek et al. 2012). And it conceals inconclusive findings, thereby preventing researchers from avoiding such methods in the future. Through these effects, HARKing distorts the evidence published in the scientific literature, compromising evidence synthesis, and scientific and medical progress.

Besides scientific and economic costs, HARKing may have additional ethical costs, if animals are used for inconclusive research or if patients in clinical trials are put at risk. A potential solution to prevent HARKing is the pre-registration of study protocols, where researchers define the hypothesis and analysis plan before a study is performed (Munafò et al. 2017; Nosek et al. 2018). Such practice compels scientists to keep with their predictions and allows for transparent reporting of what “worked” in a study and what did not. Pre-registration and more transparent reporting can aid identification of valid and reproducible test measures, advance self-correction in scientific research, and promote responsible research.

## Publication bibliography

Begley, C. Glenn; Ioannidis, John P. A. (2015): Reproducibility in science: improving the standard for basic and preclinical research. In Circulation research 116 (1), pp. 116–126. DOI: 10.1161/CIRCRESAHA.114.303819.

Bespalov, Anton; Steckler, Thomas (2021): Pharmacology of Anxiety or Pharmacology of Elevated Plus Maze? In Biological psychiatry. DOI: 10.1016/j.biopsych.2020.11.026.

Bishop, Dorothy (2019): Rein in the four horsemen of irreproducibility. In Nature 568 (435).

Ennaceur, Abdelkader; Chazot Paul L. (2016): Preclinical animal anxiety research - flaws and prejudices. In Pharmacology research & perspectives 4 (2), e00223. DOI: 10.1002/prp2.223.

Fraser, Hannah; Parker, Tim; Nakagawa, Shinichi; Barnett, Ashley; Fidler, Fiona (2018): Questionable research practices in ecology and evolution. In PloS one 13 (7), e0200303. DOI: 10.1371/journal.pone.0200303.

Kerr, Norbert L. (1998): HARKing: Hypothesising After the Results are Known. In Personality and Social Psychology Review 2 (3), pp. 196–217.

Munafò Marcus R.; Nosek Brian A.; Bishop Dorothy V. M.; Button Katherine S.; Chambers Christopher D.; Du Sert, Nathalie Percie et al. (2017): A manifesto for reproducible science. In Nature human behaviour 1, p. 21. DOI: 10.1038/s41562-016-0021.

Murphy, Kevin R.; Aguinis, Herman (2019): HARKing: How Badly Can Cherry-Picking and Question Trolling Produce Bias in Published Results? In J Bus Psychol 34 (1), pp. 1–17. DOI: 10.1007/s10869-017-9524-7.

Nosek, Brian A.; Ebersole Charles R.; DeHaven, Alexander C.; Mellor David T. (2018): The preregistration revolution. In Proceedings of the National Academy of Sciences of the United States of America 115 (11), pp. 2600–2606. DOI: 10.1073/pnas.1708274114.

Nosek, Brian A.; Spies Jeffrey R.; Motyl, Matt (2012): Scientific Utopia: II. Restructuring Incentives and Practices to Promote Truth Over Publishability. In Perspectives on psychological science : a journal of the Association for Psychological Science 7 (6), pp. 615–631. DOI: 10.1177/1745691612459058.

Prosperi, Mattia; Bian, Jiang; Buchan Iain E.; Koopman James S.; Sperrin, Matthew; Wang, Mo (2019): Raiders of the lost HARK: a reproducible inference framework for big data science. In Palgrave Commun 5 (1). DOI: 10.1057/s41599-019-0340-8.

Rosso, Marianna; Wirz, Robin; Loretan Ariane Vera; Sutter Nicole Alessandra; Pereira da Cunha, Charlène Tatiana; Jaric, Ivana et al. (2021): Reliability of Mouse Behavioural Tests of Anxiety: a Systematic Review and Meta-Analysis on the Effects of Anxiolytics. In bioRxiv : the preprint server for biology. DOI: 10.1101/2021.07.28.454267.

Rubin, Mark (2017): When Does HARKing Hurt? Identifying When Different Types of Undisclosed Post Hoc Hypothesizing Harm Scientific Progress. In Review of General Psychology 21 (4), pp. 308–320. DOI: 10.1037/gpr0000128.

Rubin, Mark (2020): The Costs of HARKing. In The British Journal for the Philosophy of Science, p. 0. DOI: 10.1093/bjps/axz050.

Wicherts, Jelte M.; Veldkamp Coosje L. S.; Augusteijn Hilde E. M.; Bakker, Marjan; van Aert Robbie C. M.; van Assen Marcel A. L. M. (2016): Degrees of Freedom in Planning, Running, Analyzing, and Reporting Psychological Studies: A Checklist to Avoid p-Hacking. In Frontiers in psychology 7, p. 1832. DOI: 10.3389/fpsyg.2016.01832.

